# Human genomics of acute liver failure due to hepatitis B virus infection: an exome sequencing study in liver transplant recipients

**DOI:** 10.1101/320374

**Authors:** Samira Asgari, Nimisha Chaturvedi, Petar Scepanovic, Christian Hammer, Nasser Semmo, Emiliano Giostra, Beat Müllhaupt, Peter Angus, Alexander J Thompson, Darius Moradpour, Jacques Fellay

## Abstract

Acute liver failure (ALF) or fulminant hepatitis is a rare, yet severe outcome of infection with hepatitis B virus (HBV) that carries a high mortality rate. The occurrence of a life-threatening condition upon infection with a prevalent virus in individuals without known risk factors is suggestive of pathogen-specific immune dysregulation. In the absence of established differences in HBV virulence, we hypothesized that ALF upon primary infection with HBV could be due to rare deleterious variants in the human genome. To search for such variants, we performed exome sequencing in 21 previously healthy adults who required liver transplantation upon fulminant HBV infection and 172 controls that were positive for anti-HBc and anti-HBs antibodies but had no clinical history of jaundice or liver disease. After a series of hypothesis-driven filtering steps, we searched for putatively pathogenic variants that were significantly associated with case-control status. We did not find any causal variant or gene, a result that does not support the hypothesis of a shared monogenic basis for human susceptibility to HBV-related ALF in adults. This study represents a first attempt at deciphering the human genetic contribution to the most severe clinical presentation of acute HBV infection in previously healthy individuals.

**Author Summary:** Infection with hepatitis B virus (HBV) is very common and causes a variety of liver diseases including acute and chronic hepatitis, cirrhosis and liver carcinoma. Acute HBV infection is often asymptomatic, still about 1% of newly infected people develop a rapid and severe disease known as acute liver failure or fulminant hepatitis. Acute liver failure has a high mortality rate and is an indication for urgent liver transplantation. It is not clear why some people, who are otherwise healthy, develop such severe symptoms upon infection with a common pathogen. Here, we hypothesized that rare DNA variants in the human genome could contribute to this unusual susceptibility. We sequenced the exome (i.e. the regions of the genome that encode the proteins) of 21 previously healthy adults who required liver transplantation upon fulminant HBV infection and searched for rare genetic variants that could explain the clinical presentation. We did not identify any variant that could be convincingly linked to the extreme susceptibility to HBV observed in the study participants. This suggests that HBV-induced acute liver failure is more likely to result from the combined influence of multiple genetic and environmental factors.

## Introduction

Hepatitis B virus (HBV) is a common human pathogen that attacks the liver and can cause both acute and chronic disease. There is high inter-individual variability in the clinical presentation of HBV infection, which ranges from self-limited to fulminant acute disease, and from mild chronic hepatitis to liver cirrhosis and hepatocellular carcinoma [1]. Differences in viral or environmental factors only explain a fraction of this variability [2–5]. Previous studies have identified some human genetic factors that play a modulating role in the clinical course of HBV infection [6,7]. However, our understanding of host genetic influences on the disease is still very limited.

Fulminant hepatitis or acute liver failure (ALF) is defined as the rapid development of liver injury leading to severe impairment of the synthetic capacity and to hepatic encephalopathy in patients without previous liver disease [8,9]. ALF due to HBV infection, or fulminant hepatitis B, is observed in less than 0.1% of infected individuals but carries a high mortality and is an indication for urgent liver transplantation [10–15].

Such an unusual clinical presentation fits the definition of an extreme phenotype. Electing patients with extreme phenotype increases the power to detect causal gene as variants as these patients are more likely to carry alleles with profound functional consequences that are otherwise very rare in the population, due to purifying selection [16–20]. In this study, we used exome sequencing and statistical analysis in a cohort of 21 cases and 172 controls to search for human genetic variants conferring extreme susceptibility to HBV. Cases were previously healthy adults who required liver transplantation for fulminant hepatitis B and controls were HBV-infected adults who did not develop fulminant hepatitis (Figure 1).

**Figure 1:**
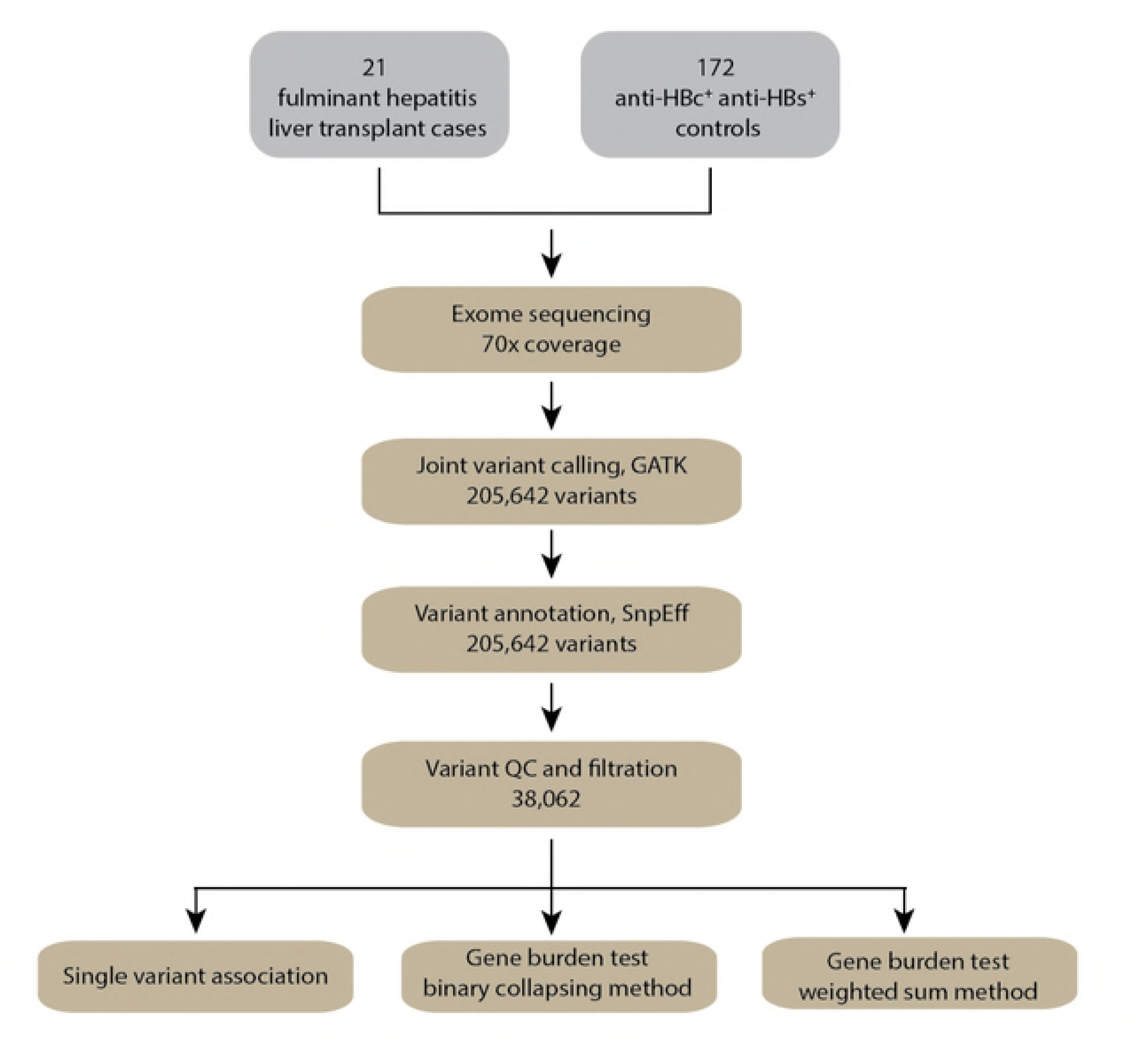
Overview of data production and data analysis pipeline. MAF: minor allele frequency, GATK: Genome Analysis Tool Kit

## Results

### Study participants

Of the 21 cases, 13 (62%) were female. The median age at transplantation was 36.5 years (range 22-58). Of 21 cases, 16 (76%) were European, four (19%) were Asian and one (5%) was African (Supplementary Figure 1).

### Exome sequencing, variant calling and variant filtering

Exome sequencing data were generated from DNA extracted from whole blood for all study participants. On average per sample, 96% of reads passing filtering criteria were unique (not marked as duplicate). Ninety-seven percent of unique reads could be aligned to the human reference genome GRCh37. The mean on-bait coverage was 73x, with 99% of target bases reaching at least 2x coverage, 97% of target bases achieving at least 10x coverage and 84% achieving at least 30x coverage. 205,642 variants were detected after GATK quality control filtering including 520 novel variants. The average transition to transversion ratio (Ti/Tv) was 2.66, and the average heterozygous to homozygous ratio was 1.5. A total of 38,062 low-frequency variants (MAF ≤ 0.05) passed filtering criteria including 31,620 rare variants (MAF ≤ 0.01, Table 1).

**Table 1:** Total number of rare (MAF ≤0.01) and rare plus low-frequency (MAF ≤0.05) variants that passes quality control and filtering criteria.

| Effect | $MAF \leq 0.01$ | $MAF \leq 0.05$ |
| --- | --- | --- |
| Inframe indel | 321 | 368 |
| Frameshift indel | 880 | 954 |
| Missense SNV | 29,549 | 35,756 |
| Splice site acceptor SNV | 152 | 177 |
| Splice site donor SNV | 177 | 200 |
| Nonsense | 541 | 607 |
| Total variants | 31,620 | 38,062 |
| Total genes | 11,595 | 12,295 |

### Single variant associate analysis

All putatively pathogenic variants were tested for association with case-control status using Fisher’s exact test. We first restricted the analysis to rare variants (MAF ≤ 0.01) and European cases only. One variant passed the Bonferroni correction threshold (p-value < 1.6e-6). The variant was a missense SNV in *IGSF3* (rs78806598, p-value=1.8e-18). Visualizing the aligned reads for this variant convinced us that this variant is called due to misalignment. This gene was excluded from our further analyses. Expanding the analysis to include the five none-European cases and the low-frequency variants did not lead to discovery of any significant associations (Supplementary Tables 1-3).

### Gene-based association analysis

11,595 genes were included in the gene burden analysis. Two different burden scores were calculated for each gene using the approached described in the methods section. Using the weighted sum method, *SLC29A1* had the lowest p-value (p-value=1.7e-5). CTSW had the lowest p-value in binary collapsing method (p-value=1.8e-5). However, none of these genes passed the Bonferroni correction threshold (p-value < 2.5e-6 for 20,000 protein coding genes, Supplementary Tables 4-5). Including the low-frequency variants (12,295 genes in total) did not change these results. The top associations including low-frequency variants were ADAM32 (p-value=1.7e-5) and PREX2 (p-value=8.7e-6) in for weighted sum method binary collapsing method respectively (Supplementary Tables 6-7). Overall, the results from the two collapsing methods and the results between rare variants (MAF ≤ 0.01) and low-frequency variants (MAF ≤ 0.05) analyses were highly concordant (Figure 2, Supplementary Figure 2). The highest correlation (*r*^2^=0.927, CI:0.924-0.929) was observed between the results of weighted sum and binary collapsing methods for rare variants. The lowest correlation (*r*^2^=0.739, CI:0.731-0.747) was observed between the results of binary collapsing method for rare and low-frequency variants (Figure 2).

**Figure 2:**
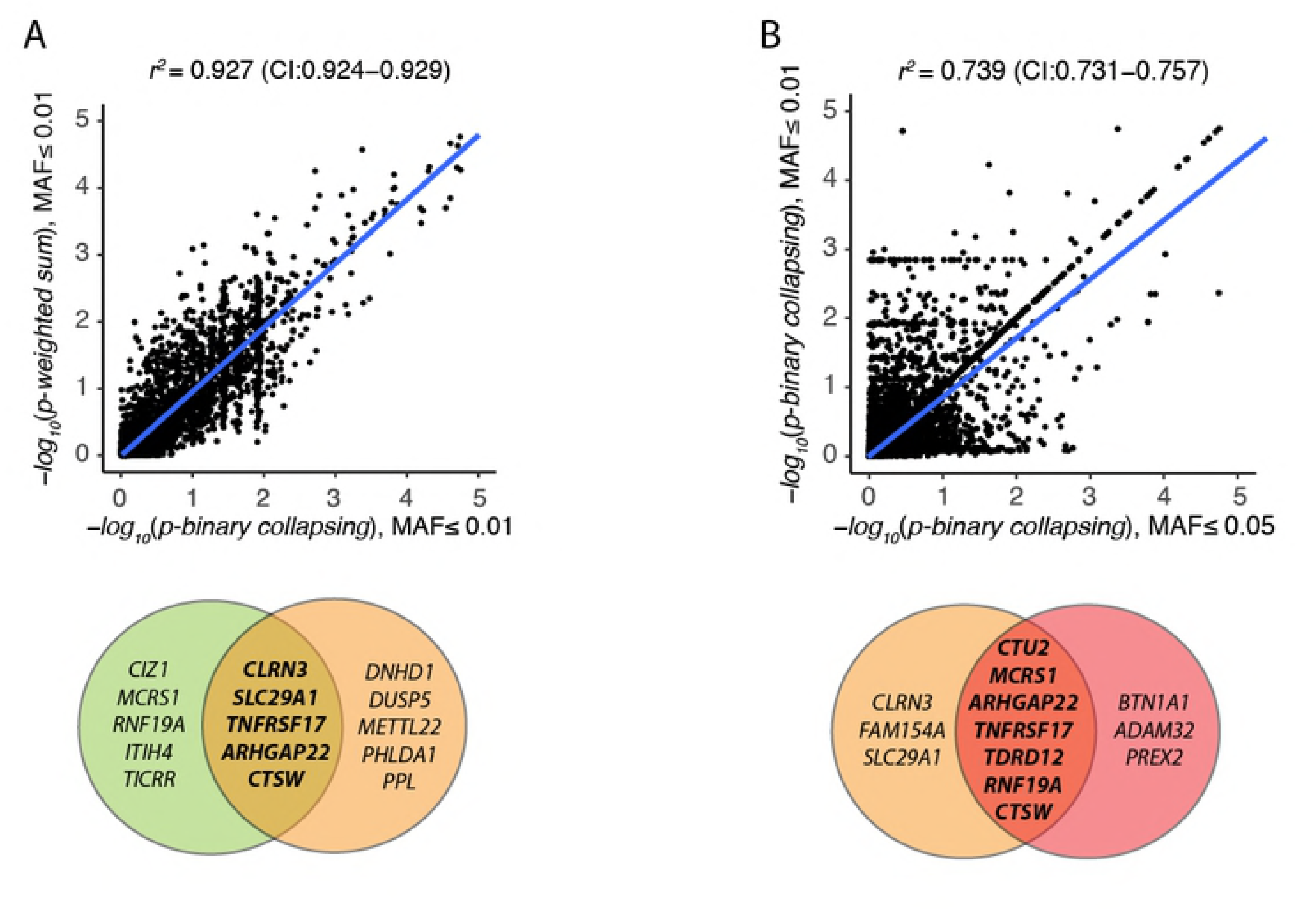
Comparison between different gene burden analysis methods and different MAF thresholds. The circles below each plot show the top ten associated genes in the two compared analyses and the number of shared genes between the two sets: A) correlation between p-value for rare variants (MAF ≤ 0.01) using weighted sum method (light green circle) and binary collapsing method (light red circle) B) correlation between p-value for binary collapsing method using MAF ≤ 0.01 (light red circle) and MAF ≤ 0.05 (dark red circle).

## Discussion

The role of human genetic factors in susceptibility to fulminant hepatitis B is poorly understood. Monogenic defects in key immune genes and pathways have been shown to cause extreme susceptibility to other common pathogens in apparently healthy individuals [7,21,22]. A prime example is herpes simplex encephalitis (HSE), the most common form of sporadic viral encephalitis in the western world, which is only observed in an extremely low fraction of people infected with type 1 herpes simplex virus (HSV-1). Children who develop HSE upon primary HSV-1 infection are not particularly susceptible to other infections, and children with other primary immunodeficiencies are not more susceptible to HSE [22]. Since 2006, multiple genetic variants have been causally linked with HSE [23–29]. Similarly, ALF only occurs in < 1/1000 of individuals after primary infection with HBV. Because this is an extremely rare clinical event, we hypothesized that it could be the first manifestation of a rare monogenic defect, resulting in pathogen-specific immune dysregulation.

We used exome sequencing to systematically search for rare, putatively pathogenic variants that could explain extreme susceptibility to HBV infection. We analyzed the genetic variants present in the exomes of 21 liver transplant recipients and compared them to 172 controls who were exposed to HBV but did not develop fulminant hepatitis B. First, we performed a single variant association analysis using Fisher’s exact test. Fisher’s exact test is a conservative test of association but guarantees type I error control for small sample sizes [30]. We found one significant association (p-value < 1.4e-6) in one gene: *IGSF3*. However, the manual inspection of the mapped reads in the region demonstrated that this variant was wrongly called, due to a mapping error. False-positive incidental findings are a major problem in small-scale exome sequencing studies [31]. Previous studies have proposed guidelines to avoid misinterpretations and erroneous reports of potential causality due to false-positive findings [32,33]. Our results show that even after applying these guidelines, it is important to ensure the quality of final findings by visualizing the mapped regions and manually verifying the quality of each variant call.

Rare variant association studies are usually underpowered. To enrich association signals and reduce the penalty of multiple testing correction, it is common to aggregate information across multiple rare variants within a region (gene, exon, sliding window, etc.) and test for the association of all variants in the region with the phenotype of interest [34]. We performed gene-based association analysis using two different aggregation methods: weighted sum collapsing and binary collapsing. Both methods assume that all the variants included in the test have the same direction of effect (increasing disease risk in our scenario) and thus are underpowered to detect disease-gene associations if variants exert their effects in opposite directions. Binary collapsing assumes that all putatively pathogenic variants have the same effect size. Weighted sum collapsing assumes that rarer variants have larger effect sizes and that the risk of disease is a function of the sum of the variant effect sizes. We did not find any genes to be significantly associated with case-control status. The p-values and the top ranked genes in both analyses were highly concordant (Figure 2, Supplementary Figure 2). The high correlation between the p-values of weighted sum and binary collapsing methods suggests that most individuals carry only one putatively pathogenic variant per gene. This implies that larger sample sizes or linkage studies in families with multiple affected individuals will be needed to increase statistical power for detecting potential associations between rare variants and HBV-related ALF.

We did not identify any genetic variant conferring monogenic susceptibility to fulminant hepatitis B in adults. Our results suggest that ALF upon primary infection with HBV is likely to be multifactorial. This conclusion is in line with a previous exome sequencing study of fulminant hepatitis A, which also failed to find any convincing casual gene or genetic variant [35]. Our failure to detect a Mendelian cause for fulminant hepatitis B, despite previous success for comparable phenotypes, could be due to a number of factors and limitations of our study: 1-The severe liver injury observed in patients with fulminant hepatitis B can be due to opposite pathogenic mechanisms: an inefficient innate immune response, which is unable to prevent viral replication, activate the adaptive immune system and clear the virus; and an over-activation of innate immune signaling pathways leading to cytokine storm and uncontrolled inflammation [36–38]. This implies that genetic variants with opposite effects (e.g. gain-of-function and loss-of-function variants in the same gene or pathway) could contribute synergistically to the disease. Such a genetic architecture would be extremely difficult to identify. 2-Our study was performed in adults, while most previous examples come from pediatric studies. A previous twin study has shown that the estimated heritability of many immune parameters decreases with age, suggesting that the cumulative influence of environmental exposures alters the role of human genetics in susceptibility to infectious diseases in older patients [39]. 3-Due to our recruitment criteria, we did not have access to information about the viral genome. HBV genetic variation has previously been shown to be associated with disease severity and infection outcome. For example, mutations in the pre-core and core protein genes or X region of the HBV genome, which encode the HBx protein, were associated with severity of chronic hepatitis [40,41], and mutations in HBV genotypes C and D associated with liver cirrhosis and progression to hepatocellular carcinoma [42]. The inclusion of viral genome information would allow for the stratification of patients based on known HBV mutations, thus increasing the signal-to-noise ratio in the human genetic analyses.

This study represents the first attempt at identifying human genetic variants involved in the pathogenesis of fulminant hepatitis B in previously healthy individuals. The absence of any conclusive finding indicates that ALF due to primary HBV infection is unlikely to be the result of a single monogenic disorder, and that a more complex genetic architecture is probably involved, intermixed with viral and environmental factors. Going forward, studies that aim at identifying the genetic causes of fulminant hepatitis B will need to include more patients and to better characterize them at the molecular level (e.g. to stratify them based on specific immune activation markers measured during acute disease). To obtain a more complete description of human genetic variation, full genome sequencing would be preferable, which will allow the exploration of non-coding variants, large structural variants and exonic variants that are not well-covered by current exome capture methods. Finally, a parallel evaluation of the viral genome and of any potentially interfering factor will be necessary, as individual susceptibility to HBV is the result of a complex interplay between host, pathogen and environment.

## Methods

### Ethics statement

The study was approved by the responsible institutional Human Research Ethics Committees in Switzerland and Australia. Each study participant provided written informed consent for genetic testing.

### Study participants

Twenty-one liver transplant recipients who developed ALF due to fulminant HBV infection were recruited in the transplantation units of the University Hospitals of Lausanne, Zurich, Bern, Geneva, and Melbourne. Patients with fulminant hepatitis B due to reactivation after withdrawal of anti-HBV drugs and patients with pre-existing liver diseases, known immune deficiency or other chronic conditions were excluded. The following demographic and clinical information were collected: age at transplantation date, gender, and ethnicity. For each study participant, we obtained 3ml of blood in EDTA vacutainer tubes and 2,5ml blood in PAXgene blood RNA tubes. Samples were immediately frozen at −70°C, and then shipped and analyzed in batch.

### Control population

One hundred seventy-two controls were selected from our in-house database of exome-sequenced individuals. They were adults of European ancestry, who were positive for anti-HBc and anti-HBs antibodies, but had no clinical history of jaundice or liver disease. The controls were HBV-eliminated at the time of blood collection for exome-sequencing.

### DNA sequencing and alignment

Genomic DNA was extracted from whole blood using QIAgen DNeasy Blood and Tissue kit. Cluster generation was performed using Illumina TruSeq PE Cluster Kit v5 reagents. Libraries were sequenced as 100-basepair long, paired-end reads on Illumina HiSeq 2500 using TruSeq SBS Kit v5 reagents. Sequencing reads were processed using CASAVA v1.82, and aligned to the human reference genome hg19 using BWA [43,44] version 0.6.2. PCR duplicates were removed using Picard 1.27-1 (http://picard.sourceforge.net/). We used Samtools [45] Visualization of aligned reads.

### Variant calling

We used Genome Analysis Toolkit (GATK) [46,47] version 3.1-1 to call single nucleotide variants (SNVs) and small insertion and deletions (indels) from duplicate-marked bam files. We used HaplotypeCaller for multi-sample variant calling on all samples following GATK best practice.

### Variant effect prediction, frequency estimation and filtering

We used SnpEff [48] version 4.3T to predict the functional impact of variants. As a single variant can have several predicted effects, we only considered the most severe effect for each variant according to SnpEff order of impact severity. We used genome aggregation database (gnomAD) to assign minor allele frequency (MAF) to variants (gnomAD, includes 123,136 exome sequences and 15,496 whole-genome sequences) [49]. For variants that were not present in gnomAD were assigned MAF=1-e8 to avoid having −log(0) in the following burden analysis. Only biallelic variants that were flagged as PASS by GATK, and were called in all cases and all controls were included in the analysis. Known polymorphic genes and genes in noisy alignment regions were excluded from the analysis [31–33]. We restricted all the downstream analyses to protein modifying variants (missense, inframe indels, frame-shift indels, splice-site disrupting, nonsense). All analyses were done on both rare (MAF ≤ 0.01) and low-frequency (MAF ≤ 0.05) variants. We refer to variants that passed above filtering criteria as putatively pathogenic variants.

### Single variant association tests

We used Fisher’s exact test to look for association of single variants with case-control status. Each variant was given an allele count based on the number of alternate alleles *G_ij_* ∈ {0,1,2}, where *G_ij_* is the genotype of variant *j* in individual *i*. We summarized the reference and alternate allele counts for cases and controls, into 2×2 contingency tables. These tables were analyzed using one-tailed Fisher’s exact test. We used Bonferroni correction to correct for multiple testing.

### Gene burden association tests

Gene burden test was performed using GMMAT [50] version 0.7-1, a generalized linear mixed model framework as follows:

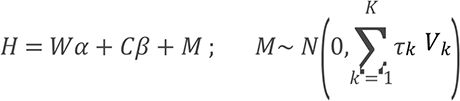

Where *H* is an *n*-vector of case-control status for *n* individuals, *W* is an n-vector of gender covariate, and *C* is an *n*-vector of the gene burden scores. M is an n-vector of random effects. is the variance component parametes and *V_k_* are known *n × n* matrices. We ran this model using an *n × n* kinship coefficients matrix calculated using PC-Relate [51]. We used Bonferroni correction to correct for multiple testing. To calculate the gene burden scores, we used two different methods:

#### i) Binary collapsing method

Each gene was given a burden score of zero if no putatively pathogenic variant was present in the gene and a gene burden score of one otherwise:

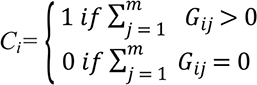

Where *G_ij_ ∈* {0,1,2} is the genotype of variant *j* in individual *i*, and *C_i_* is the gene burden score for individual *i*. This approach is based on the Cohort Allelic Sum Test (CAST) method [52].

#### ii) Weighted sum collapsing method

First, each gene was given a burden score as follows:

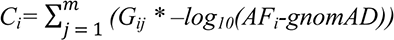

Where *j* ∈ {0,1,2,…,*m*} is the mth variant per gene, and *G_ij_* ∈ {0,1,2} is the genotype of variant *j* in individual *i, AF_i_*-*gnomAD* is the minor allele frequency of *j* in gnomAD, and *C_i_* is the gene burden score for individual *i*. This approach is based on the Madsen and Browning weighted sum method [53].

## Acknowledgements

Sequencing was performed at the Lausanne Genomic Technologies Facility of the University of Lausanne. The computations were performed at the Vital-IT (http://www.vital-it.ch) Center for high-performance computing of the SIB Swiss Institute of Bioinformatics. We would like to thank the study participants, as well as the study nurses, physicians and laboratories who participated in the recruitment.

## Supplementary Tables legends

Supplementary Tables 1-3: **Fisher’s exact test results for single variant association analysis** for: 1-Rare variants (MAF ≤ 0.01), 16 European cases and 172 controls, 2-Rare variants (MAF ≤ 0.01), all 21 cases and 172 controls, 3-Low-frequency variants (MAF ≤ 0.05), all 21 cases and 172 controls. Column names: chromosome, position, reference allele, alternate allele, variant ID, gene, number of putatively pathogenic alleles in cases, number of putatively pathogenic alleles in controls, number of non-putatively pathogenic alleles in cases, number of non-putatively pathogenic alleles in controls

Supplementary Tables 4-7: **Gene burden association results** for: 4-Rare variants (MAF ≤ 0.01) and weighted sum method, 5-Rare variants (MAF ≤ 0.01) and binary collapsing method, 6-Rare variants (MAF ≤ 0.05) and weighted sum method, 7-Rare variants (MAF ≤ 0.05) and binary collapsing method. Column names: gene, score, variance, p-value

## Supplementary Figures legends

Supplementary Figure 1: **Principal component analysis (PAC)** A-B) PCA analysis for 21 cases, 172 controls and continental populations from 1000 genomes project, C-D) PCA analysis for 21 cases and 172 controls.

Supplementary Figure 2: **Comparison between different gene burden analysis methods and different MAF thresholds.** The circles below each plot show the top ten associated genes in the two compared analyses and the number of shared genes between the two sets: A) correlation between p-value for low-frequency variants (MAF ≤ 0.05) using weighted sum method (dark green circle) and binary collapsing method (dark red circle). B) correlation between p-value for weighted sum method using MAF ≤ 0.01 (light green circle) and MAF ≤ 0.05 (dark green circle).

